# High-fidelity, high-spatial-resolution diffusion MRI of the ex-vivo whole human brain on the 3T Connectom scanner using structured low-rank EPI ghost correction

**DOI:** 10.1101/2021.08.01.454635

**Authors:** Gabriel Ramos-Llordén, Rodrigo A. Lobos, Tae Hyung Kim, Qiyuan Tian, Thomas Witzel, Hong-Hsi Lee, Alina Scholz, Boris Keil, Anastasia Yendiki, Berkin Bilgiç, Justin P. Haldar, Susie Y. Huang

**Author notes:** Correspondence to: Gabriel Ramos-Llordén, PhD.

## Abstract

Diffusion MRI (dMRI) of whole, intact, fixed postmortem human brain at high spatial resolution serves as key bridging technology for 3D mapping of structural connectivity and tissue microstructure at the mesoscopic scale. Ex vivo dMRI offers superior spatial resolution compared to in vivo dMRI but comes with its own technical challenges due to the significantly reduced T2 relaxation times and decreased diffusivity incurred by tissue fixation. The altered physical properties of fixed tissue necessitate the use of alternative acquisition strategies to preserve SNR and achieve sufficient diffusion weighting. Multi-shot or segmented 3D echo planar imaging (EPI) sequences have been used to shorten echo times (TEs) with reduced distortions from field inhomogeneity and eddy currents on small-bore MR scanners and have been adopted for high b-value dMRI of ex vivo whole human brain specimens.

The advent of stronger gradients on human MRI scanners has led to improved image quality and a wider range of diffusion-encoding parameters for dMRI but at the cost of more severe eddy currents that result in spatial and temporal variations in the background magnetic field, which cannot be corrected for using standard vendor-provided ghost correction solutions. In this work, we show that conventional ghost correction techniques based on navigators and linear phase correction may be insufficient for EPI sequences using strong diffusion-sensitizing gradients in ex vivo dMRI experiments, resulting in orientationally biased dMRI estimates. This previously unreported problem is a critical roadblock in any effort to leverage scanners with ultra-high gradients for high-precision mapping of human neuroanatomy at the mesoscopic scale. We propose an advanced reconstruction method based on structured low-rank matrix modeling that reduces the ghosting substantially. We show that this method leads to more accurate and reliable dMRI metrics, as exemplified by diffusion tensor imaging and high angular diffusion imaging analyses in distributed neuroanatomical areas of fixed whole human brain specimens. Our findings advocate for the use of advanced reconstruction techniques for recovering unbiased metrics from ex vivo dMRI acquisitions and represent a crucial step toward making full use of strong diffusion-encoding gradients for neuroscientific studies seeking to study brain structure at multiple spatial scales.

## 1. Introduction

Ex vivo diffusion MRI (dMRI) serves as a key bridging modality for probing structural connectivity of the human brain at the mesoscopic scale (Roebroeck, Miller, & Aggarwal, 2019). Diffusion MRI measurements performed on post-mortem fixed human brain tissue can achieve much higher spatial resolution (~a few hundred microns isotropic voxel size) than is achievable in vivo (~1-2 mm isotropic voxel size) while accessing a larger field of view, up to the whole brain level using large-bore MRI scanners, compared to most microscopic imaging techniques (Miller, McNab, Jbabdi, & Douaud, 2012) (Miller, et al., 2011) (McNab, et al., 2013) (McNab, et al., 2009) (Fritz, et al., 2019). Ex vivo dMRI also provides an excellent testbed for validating in vivo dMRI measures of structural connectivity, including subcortical and intracortical fiber orientations, and tissue microstructure (Roebroeck, Miller, & Aggarwal, 2019) (Miller, et al., 2011) (Leuze, et al., 2014) (Aggarwal, Nauen, Troncoso, & Mori, 2015) (Yendiki, Aggarwal, Axer, Howard, & Haber, 2021) (Kleinnijenhuis, et al., 2013) (Assaf, 2019) (Dinse, et al., 2015) (Beaujoin, et al., 2018) (Augustinack, et al., 2010).

The acquisition of high spatial resolution ex vivo dMRI data in whole human brain specimens poses significant technical challenges, which are driven by the altered chemical properties incurred by tissue fixation. Perhaps the most striking differences between ex vivo and in vivo dMRI are the substantially reduced T2 relaxation time (e.g., ~45-55 ms in ex vivo human white matter compared to ~70 ms in vivo (Deoni, 2011)) and diffusivity of fixed tissue (e.g., 0.8 × 10^−4^ *mm*^2^ / *s* in fixed white matter) (Shepherd, Thelwall, Stanisz, & Blackband, 2009) (McNab, et al., 2009) (Miller, McNab, Jbabdi, & Douaud, 2012) (Foxley, et al., 2014). The shortened T2 and reduced diffusivity can be offset through the use of high gradient strengths for stronger and more efficient diffusion-encoding, longer acquisition times, highly sensitive radiofrequency coils, and optimized image-encoding strategies (Roebroeck, Miller, & Aggarwal, 2019) (Dyrby, et al., 2011) (Calamante, et al., 2012) (Eichner, et al., 2020a) (Eichner, et al., 2020b). For example, these strategies have been demonstrated in ex vivo dMRI measurements performed on human and non-human brains using small-bore MR scanners through the use of spin-echo sequences and 3D encoding (D’Arceuil, Westmoreland, & de Crespigny, 2007) (Roebroeck, et al., 2008) (Calabrese, et al., 2015). Such strategies have been less explored on large-bore human MRI scanners, which have been traditionally limited to gradient strengths <100 mT/m. The introduction of gradient coils capable of attaining maximum gradient amplitudes up to 300 mT/m in human MRI scanners (Setsompop, et al., 2013) (McNab, et al., 2013) (Jones, et al., 2018) and dedicated, highly parallel radiofrequency arrays for ex vivo brain imaging (Scholz, et al., 2021) have paved the way to achieving high *b*-value, high sensitivity ex vivo dMRI of whole human brain specimens.

Nevertheless, the use of strong diffusion-sensitizing gradients presents challenges to the image encoding, which must be accounted for in order to avoid image artifacts resulting in spurious estimates of dMRI measures. The most common image readout strategy used in in vivo dMRI, single-shot EPI (ss-EPI), is not well-suited for ex vivo dMRI due to the shorter T2 relaxation times in fixed tissue. The need for shorter TE in ex vivo dMRI measurements directly contradicts the use of lengthy echo trains in ss-EPI, which are further prolonged when using large matrix sizes for achieving sub-millimeter spatial resolution. Therefore, ex vivo dMRI experiments often employ multi-shot or segmented EPI readout strategies, which combine interleaved k-space trajectories from multiple excitations using shorter echo trains to reduce the overall TE. Previous studies have used 3D multi-shot EPI-based acquisitions to achieve dMRI of whole post-mortem human brain specimens at submillimeter isotropic spatial resolution on 3T human scanners (Miller, et al., 2011) (McNab, et al., 2013). With the advent of higher gradient strength human MRI scanners, the eddy currents that are generated with the rapid switching of strong diffusion-sensitizing gradients create temporal and spatial variations of the magnetic field that are more severe than those seen in conventional clinical MRI scanners. These time- and spatially-varying magnetic fields introduce systematic differences in the interleaved k-space lines acquired with alternating gradient polarities and at different shots or excitations. If not properly accounted for, profound Nyquist ghosting appears in the reconstructed images that cannot be fixed using conventional linear phase correction strategies, thereby motivating the need for more advanced image reconstruction techniques.

In this work, we address the ghosting artifacts incurred by gradient system-induced eddy currents in high spatial resolution whole brain ex vivo dMRI acquired with strong diffusion-sensitizing gradients on the 3T Connectom scanner. We show that the ghosting artifacts observed in dMRI experiments using gradient strengths approaching the maximum gradient amplitude of 300 mT/m and 3D multi-shot EPI readout cannot be mitigated with conventional ghost correction approaches, i.e., linear phase correction with one-dimensional navigators, due to simplistic phase modelling assumptions that do not apply in this high-gradient strength regime. Furthermore, the prevailing ghosting artifacts interfere with the estimation of scalar and orientational dMRI metrics such as those derived from diffusion tensor imaging (DTI). We then propose an alternative method for ghost correction that relies on advanced reconstruction with concepts from structured low-rank matrix (SLM) modeling theory (Lobos, Kim, W.S, & Haldar, 2018) (Mani, Jacob, Kelley, & Magnotta, 2017) (Lee, Jin, & Ye, 2016) (Shin, et al., 2014).

We show that this advanced reconstruction approach reduces ghosting artifacts substantially, without introducing additional artifacts or apparent penalties in SNR. The SLM-based EPI correction method is first validated in an isotropic diffusion phantom and then applied to dMRI experiments performed in a fixed whole human brain specimen acquired at 0.8 mm isotropic resolution and *b*-values up to 10 000 s/mm^2^. We compare the estimation of commonly used dMRI metrics, including DTI measures and fiber orientation distribution functions (fODFs), from images reconstructed using conventional linear phase correction and SLM-based EPI ghost correction and demonstrate the improved mapping of dMRI metrics in key neuroanatomical areas distributed across the whole brain.

## 2. Material and Methods

### 2.1 EPI ghosting artifacts from strong diffusion-sensitizing gradients

The work presented here makes use of a 3D multi-shot or segmented EPI diffusion-weighted spin echo sequence (Miller, et al., 2011) (McNab, et al., 2013). In a multi-shot EPI sequence, k-space is sampled using multiple excitations (shots) with complementary k-space trajectories that are later combined in an interleaved fashion (Fig 1). The 3D multi-shot EPI sequences commonly used for ex vivo dMRI employ a 3D k-space readout, which enables efficient whole brain coverage and boosts SNR over the 2D readouts commonly used for in vivo dMRI.

**Fig 1.**
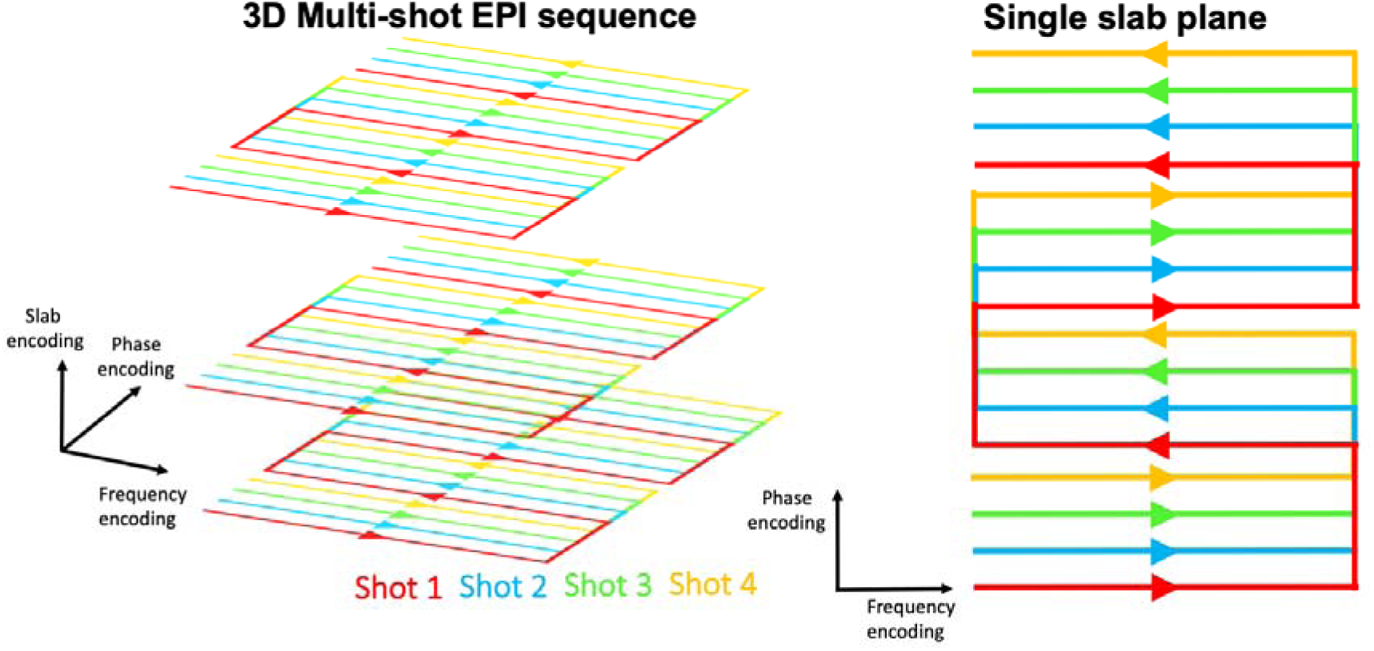
The 3D Multi-shot EPI sequence used in this work. Each 2D k-space plane (fixed slab encoding axis) is acquired with a 2D multi-shot EPI sequence, where k-space is divided into four segments: red, blue, green, and dark blue, each one acquired at consecutive TR’s. Within each segment, the k-space plane is under-sampled by a factor of four. Once all segments are interleaved, the k-space data is fully sampled

Due to their shorter effective echo spacings, multi-shot or segmented k-space trajectories are less prone to geometric distortions, relaxation-related blurring and SNR loss than ss-EPI (Roebroeck, Miller, & Aggarwal, 2019). Nevertheless, some of the intrinsic drawbacks of EPI are still present, including its sensitivity to temporal and spatial variations of the main magnetic field (B_0_). Deviations of the main magnetic field from the nominal B_0_ field and hence, from the prescribed k-space trajectory, can arise from susceptibility differences, field inhomogeneities, chemical shift, as well as numerous hardware imperfections (Reeder, Atalar, & E.R., 1997) and eddy currents (Buonocore & Gao, 1997). Eddy currents are induced in conducting structures due to temporally and spatially varying magnetic fields (Boesch, Gruetter, & Martin, 1991). In dMRI, the rapid switching of the diffusion-encoding gradients is the main source of eddy currents. Eddy currents produce magnetic field gradients that persist after the diffusion-sensitizing gradients are turned off, with a finite lifetime that overlaps with the image readout (Le Bihan, Poupon, Amadon, & Lethimonnier, 2006). Aside from geometric distortions, which can be corrected with post-processing tools such as ‘*topup*’ and ‘*eddy*’ from the FSL software (Andersson & Sotiropoulos, 2016), an unwanted phase modulation appears in the k-space data. If not properly corrected, Nyquist-ghosting artifacts emerge in the reconstructed images. The focus of this paper is on these phase modulations incurred by eddy currents.

The literature of EPI ghosting correction is rich and a comprehensive review of it is beyond the scope of this work. However, it is instructive to place the proposed methodology into one of the two main categories of ghost correction methods: phase correction approaches that rely on navigator data (Hu & Le, 1996) (Chen & Wyrwicz, 2004) (Bruder, Fischer, Reinfelder, & Schmitt) (Hennel, 1998), and other techniques that make fewer assumptions on the model but instead treat the ghosting problem as an inverse reconstruction problem in which techniques adapted from parallel imaging can be used (Hoge & Polimeni, 2016) (Lee, Jin, & Ye, 2016) (Xie, Lyu, Liu, Feng, & Wu, 2018) (Lobos, Kim, W.S, & Haldar, 2018) (Lobos, et al., 2021). Note that in this classification we are omitting other techniques that focus on mitigating eddy current effects in the acquisition, e.g., gradient shielding (Mansfield & Chapman, 1986), pre-emphasis or dual spin-echo sequences (Reese, Heid, Weisskoff, & Wedeen, 2003), and other hardware solutions such as higher-order field monitoring (Wilm, Barmet, Pavan, & Pruessmann, 2011). The methodology presented here can be readily applied in combination to those techniques. We further elaborate on this matter in the Discussion section.

Phase correction approaches assume low-dimensional models for the phase modulation between k-space lines of different polarities and shots. The parameters of the model are estimated from calibration scans (Buonocore & Gao, 1997). While these methods are yet widely used in common dMRI acquisitions with conventional gradient strengths (40 mT/m), we show here that their applicability to the regime of strong diffusion-encoding gradients is severely limited. As eddy currents increase with the gradient strength, they turn out to be the main source of ghosting in the high-gradient strength regime, leading to phase modulations that are not fully captured by the simple models that phase navigator-based methods use (Lobos, Kim, W.S, & Haldar, 2018). The methodology that is applied here falls under the second category of methods and is described below.

### 2.2 EPI ghosting reduction with SLM-based reconstruction

We will start with the assumption that the 3D k-space data have been inverse Fourier transformed along the slab direction, which is fully sampled. If so, the image reconstruction problem can be treated in a slice-wise fashion. For a given slab/slice and diffusion direction, let 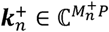 be a complex vector which contains the multi-channel (*P* is the number of channels) k-space data acquired at the n-th shot/excitation, from a total of *N_sh_* shots, comprising only k-space lines with positive readout polarity (+), e.g. considering only the even phase-encoding lines (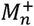 is the number of acquired k-space data points).

SSimilarly, let 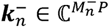 stands for the k-space dataset with the remaining k-space lines acquired with negative readout polarity (−), e.g. considering only the odd phase-encoding lines (see k-space sampling patterns in Fig 2.b). The basic premise is that both 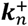 and 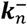 represent different images 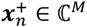 and 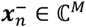 (*M* is the number of image pixels), and they are related by the following image formation model:

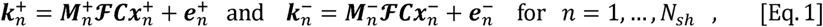

with 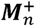 and 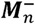 diagonal matrices (dimensions 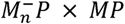 and 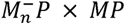), representing the under-sampling k-space masks at the *n*-th shot and positive and negative readout, respectively, matrix 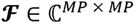 representing the 2D Fourier transform, 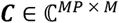 the matrix containing the coil sensitivities, and 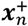 and 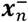 the underlying complex images from the *n*-th shot and positive and negative readouts, respectively. Finally, 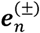 denote the measurement noise (same dimensions as 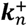 and 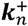, respectively). Note that as the k-space data from different shots and readout polarities have been separated, the images 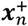 and 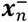 should be in principle free from Nyquist ghosting. However, reconstructing the images from 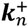 and 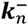 is a highly under-sampled and ill-posed problem, and conventional parallel imaging methods such as sensitivity encoding (SENSE) (Pruessmann, Weiger, Scheidegger, & Boesiger, 1999) and Generalized Autocalibrating Partially Parallel Acquisitions (GRAPPA) (Griswold, et al., 2002) methods may not provide satisfactory results at high acceleration/segmentation factors.

**Fig 2.**
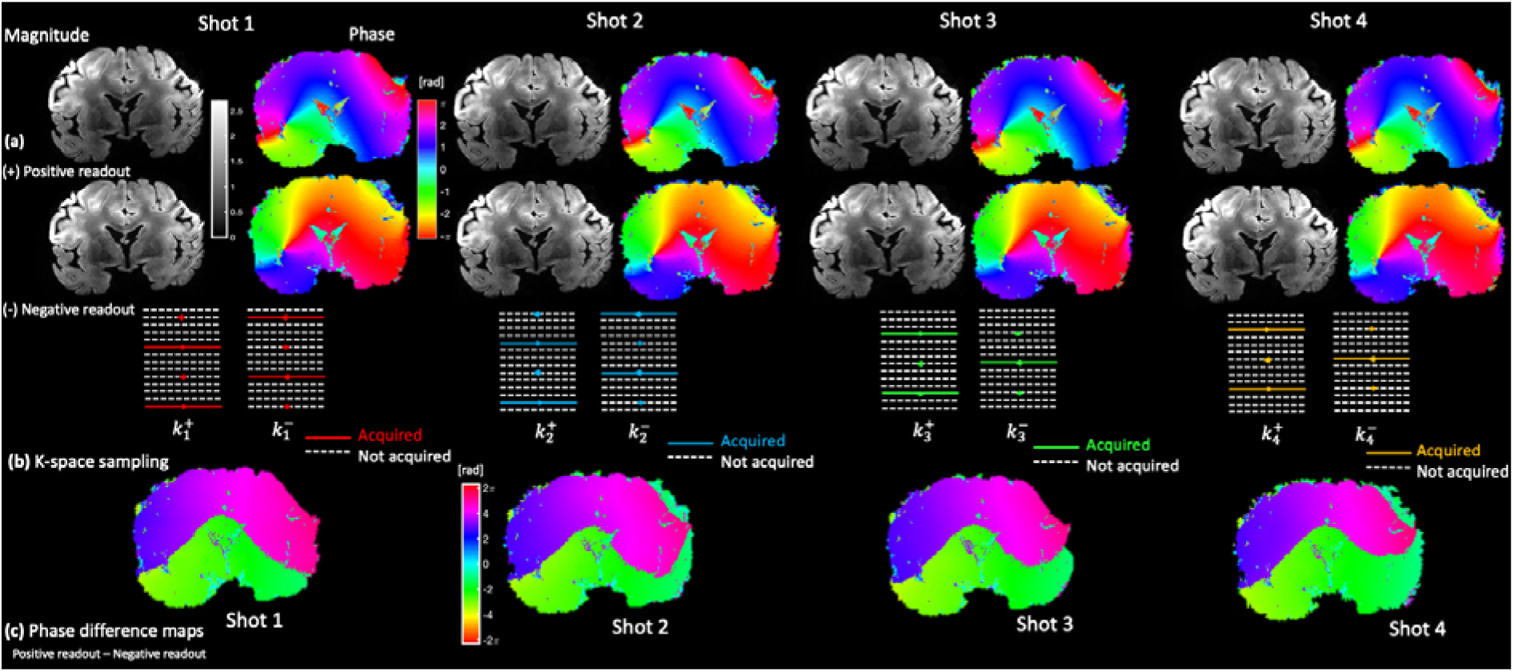
With the proposed EPI correction technique based on structured low rank-matrix modeling, a complex image (a) is reconstructed for each shot and readout polarity (positive and negative). To solve this ill-conditioned under-sampled problem, a joint regularization term (coupling the problem in a unified way) is included, which takes advantage of the linear relationships between the k-space data across shots and polarities. Next, all the magnitude images are combined with the root sum of squares technique. Coil sensitives are estimated using a separate gradient-recalled image acquisition. Note that the differences between the phase maps (c) of the reconstructed images for different polarities are highly nonlinear, underscoring the fact that traditional methods based on linear phase correction may be insufficient to reduce ghosting.

Fortunately, we can leverage the fact that the images 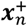 and 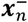 are highly correlated by including a regularization term that imposes constraints on these images. In this work, we incorporate constraints derived from the theory of SLM. SLM-based methods have been successfully applied in conventional MRI undersampling problems (Haldar J. P., 2013) (Shin, et al., 2014) (Jin, Lee, & Ye, 2015) (Kim, Setsompop, & Haldar, 2017), and recently in more problem-specific scenarios such as single-shot EPI-ghosting correction (Lee, Jin, & Ye, 2016) (Lobos, Kim, W.S, & Haldar, 2018) (Lobos, et al., 2021) (Mani, Aggarwal, Magnotta, & Jacob, 2020) or in vivo diffusion MRI multi-shot reconstruction (Mani, Jacob, Kelley, & Magnotta, 2017) (Mani, Aggarwal, Magnotta, & Jacob, 2020).

SLM-based methods exploit the fact that, if k-space data are linearly predictable, then a Hankel or Toeplitz matrix from those data is expected to have low rank (Haldar & Setsompop, 2020). Linear dependencies in the k-space data may appear due to limited image support, smooth image phase (Haldar J. P., 2013), and similarity of the magnitude across images acquired at different shots (Mani, Jacob, Kelley, & Magnotta, 2017).

In this work, we assume that intra-shot k-space linear dependencies exist, i.e. the images 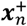 and 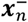 have finite image support. We also leverage inter-shot and inter-polarity k-space dependencies, that is, we assume similarity in the magnitude along polarities and shots. In this way, our approach combines the ideas of (Lobos, Kim, W.S, & Haldar, 2018) and (Mani, Jacob, Kelley, & Magnotta, 2017).

The block-Hankel matrix ***H*** which is constructed as described in (Bilgic, et al., 2019), contains k-space data of the underlying images 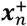 and 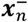, and is expected to be a low-rank matrix. To encourage ***H*** to be a low-rank matrix, we used the non-convex cost function originally presented in (Haldar J. P., 2013), and later used in (Lobos, Kim, W.S, & Haldar, 2018). This non-convex function has been shown to provide better results (Lobos, Kim, W.S, & Haldar, 2018) than other approaches used in the SLM literature, for instance the nuclear norm used in (Lee, Jin, & Ye, 2016) and (Mani, Jacob, Kelley, & Magnotta, 2017). The non-convex cost function 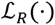 is written as:

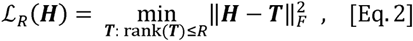

where *R* is a user-defined parameter and 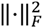 represents the Frobenius norm. This regularizer encourages matrix ***H*** to be well approximated by a matrix of rank *R* or less (Haldar J. P., 2013).

To summarize, the advanced EPI-ghosting correction framework for ex vivo dMRI consists of the following regularized optimization problem:

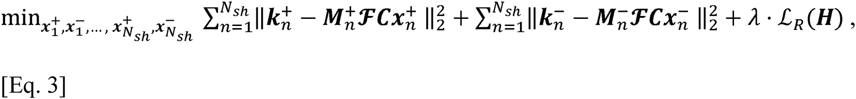

where λ is a tunable regularization parameter that controls the influence of the low-rank regularizer 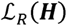 over the data fidelity term. Though the dependence has been omitted for clarity, note that the structured low-rank matrix ***H*** is a function of both 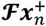 amd 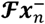, for 1,…, *N_sh_*. Coil sensitivity profiles ***C*** are estimated a priori with the ESPIRiT algorithm from a gradient echo dataset (Uecker, et al., 2014). Once reconstructed, the images are combined with the root of sum of squares technique to provide a single ghost-free reconstructed image per slice and diffusion direction.

The minimization problem of Eq. [3] is solved with a majorize-minimization (MM) approach (Haldar J. P., 2013) (Jacobson & Fessler, 2007), which is shown to decrease the cost function monotonically (Sun, P. Babu, & Palomar, 2017). Convergence of the algorithm is accelerated with the technique proposed in (Varadhan & Poland, 2008).

Three parameters should be defined by the user: the regularization parameter λ, the expected rank of the matrix ***H***, *R*, and the kernel size of the k-space neighborhoods from which ***H*** is constructed. In this work, we have not attempted to optimize the selection of the parameters according to any specific criterion. Instead, they have been chosen heuristically as those that provided the best results after exhaustive experiments. A value of λ = 2 · 10^−3^ gave the best results in our experimental validation. We observed that the performance was not very sensitive to the choice of the rank *R*, provided values between 80 and 120 were selected. We used a fixed value of 100 for all the experiments. This agrees with previous findings about the robustness of the performance of SLM methods with respect of the rank (Lobos, et al., 2021). We, however, noticed that the size of the k-space neighborhoods that are used to construct low-rank matrix, needed to be substantially higher (17 × 17) than those values reported in the literature (Bilgic, et al., 2019). A higher spatial resolution or more prominent ghosting artifacts due to the use of ultra-high gradient strengths may be one of the possible reasons to increase the size of the k-space neighborhoods. A more extensive discussion about the optimization of the algorithm is given at the end of the manuscript.

A set of reconstructed diffusion-weighted images (DWI) with the SLM-method described above, and which were acquired at 0.8-mm isotropic resolution with shots are shown in Fig 2. Note that the reconstructed magnitude images are highly similar across shots and polarities but phase images are not.

## 2.3 Experiments

### 2.3.1 Data acquisition and processing

The proposed methodology was first validated with an isotropic diffusion phantom containing 40% Polyvinylpyrrolidone (PVP) solution (Pierpaoli, Sarlls, Nevo, Basser, & Horkay, 2009). Dataset was acquired in the PVP phantom at 1 mm isotropic resolution using a *b*-value of 2000 s/mm^2^ and 24 non-collinear diffusion directions. Three initial *b*=0 images were acquired. The relevant acquisition parameters were: TE/TR = 58/3000 ms, EPI factor = 53, N_sh_ = 4, partial Fourier factor of 6/8, transverse slices, and phase-encoding in the anterior-posterior direction. Due to space constraints in the 48-channel receive coil, the phantom was scanned using a custom-built 64-channel head receive coil (Keil, et al., 2013) to accommodate the larger volume of the spherical phantom. The maximum diffusion-encoding gradient strength was 145 mT/m. Additional image acquisition parameters are given in Table 1.

**Table 1.** Acquisition parameters for the experiments on the isotropic diffusion phantom and the ex-vivo human brain.

| Parameters | Isotropic Phantom | Ex-vivo Dataset 1: Single-shell | Ex-vivo Dataset 2: Multi-shell |
| --- | --- | --- | --- |
| Resolution | 1 mm isotropic | 0.8 mm isotropic | 0.8 mm isotropic |
| TE/TR (ms) | 58 / 3000 | 79 / 500 | 60 / 500 |
| FOV (mm) | 220 x 210 x 6 | 200 x 134 x 179 | 200 x 134 x 166 |
| Matrix size<br>(read x phase x slice) | 220 x 210 x 6 | 250 x 168 x 224 | 250 x 168 x 208 |
| Number of shots | 4 | 4 | 4 |
| EPI factor | 53 | 42 | 42 |
| Effective echo spacing (ms) | 1.20 | 1.24 | 1.24 |
| Partial Fourier | 6/8 | Off | 6/8 |
| Phase encoding | A>>P | A>>P | A>>P |
| b-value(s) ($\text{s}/\text{mm}^2$) | 2 000 | 4 000 | 4 000 and 10 000 |
| Number of directions | 24 non-collinear directions | 48 non-collinear directions | 48 non-collinear directions per shell |
| b0-images (number/interleave) | 3/0 | 6/8 | 6/8 |
| Max. gradient strength (mT/m) | 145 | 180 | 153 (lower shell) and 280 (upper shell) |
| Scan time (h) | 32 min | 5 h 40 min | 10 h 42 min |

Next, a whole fixed human brain from a male who died of non-neurological causes was scanned on a dedicated high-gradient 3T MRI scanner (MAGNETOM Connectom, Siemens Healthineers) equipped with maximum gradient strength of 300 mT/m and slew rate of 200 T/m/s. The excised brain was placed in fixative (10% formaldehyde) for 90 days and transferred to paraformaldehyde-lysine-periodate solution for long-term storage. The *ex vivo* human brain was scanned with a 48-channel receive array coil specially constructed for high-sensitivity mesoscopic dMRI of whole human brain specimens (Scholz, et al., 2021). The ex vivo human brain was positioned in the dedicated whole brain ex vivo coil as shown in Fig 3. The head-first supine anatomical coordinate system (inferior-superior, left-right, anterior-posterior) in relation to the ex vivo brain is indicated in the figure. Note that with this convention, the neuroanatomical coordinates of the brain are related to the anatomical coordinate system as follows: ventral-dorsal is anterior-posterior, left-right is right-left, and rostral-caudal is inferior-superior.

**Fig 3.**
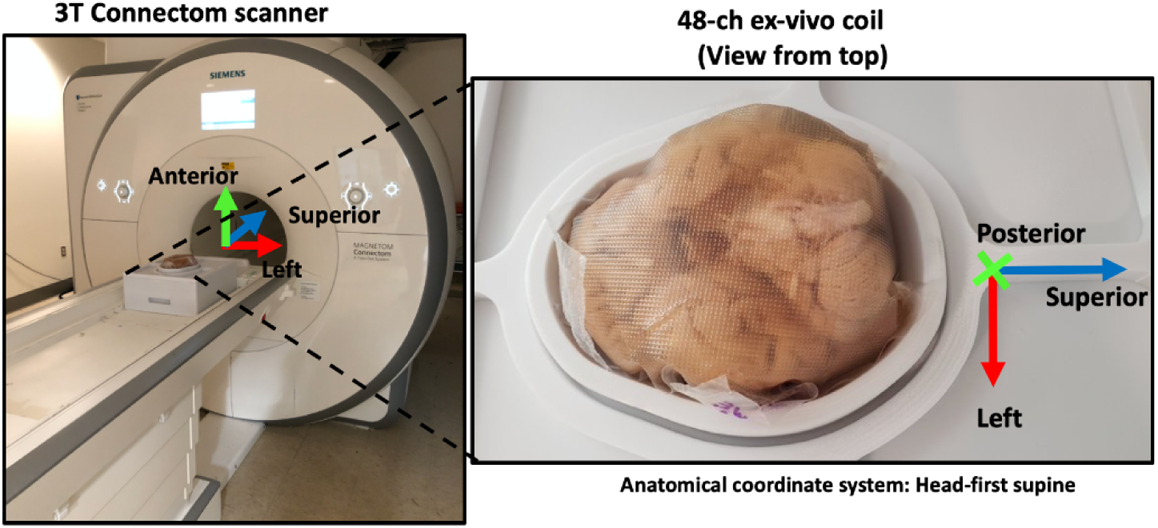
The fixed postmortem whole human brain was imaged in the 48-channel whole brain receive array coil designed for mesoscopic ex-vivo diffusion MRI. Note that in a head-first supine coordinate system, the ventral-dorsal axis of the brain corresponds to the anterior-posterior axis, the left-right axis of the brain corresponds to right-left axis, and the rostral-caudal axis of the brain is aligned with the inferior-superior anatomical axis.

The ex vivo human brain was scanned twice in separate scan sessions using different maximum gradient strengths. In the first session, we acquired a single-shell dMRI dataset at 0.8 mm isotropic spatial resolution using *b* = 4 000 s/mm^2^ and 48 non-collinear diffusion directions (maximum gradient strength of 180 mT/m, Ex-vivo Dataset 1). We employed a 3D diffusion-weighted multi-shot spin-echo sequence with the following parameters: TE/TR = 79/500 ms, EPI factor = 42, N_sh_ = 4, transverse slices, and phase-encoding in the anterior-posterior direction (ventral-dorsal direction of the brain). In this first acquisition, partial Fourier was not used to avoid introducing potential further blurring. The remainder of the relevant parameters can be found in Table 1.

The second scan session involved a multi-shell dMRI protocol acquired at the same spatial resolution as the first and *b*-values of 4 000 and 10 000 s/mm^2^ (gradient strength of 153 mT/m and 280 mT/m, respectively; Ex-vivo Dataset 2) with 48 non-collinear diffusion directions. The acquisition parameters included: TE/TR = 60/500 ms, EPI factor = 42, N_sh_ = 4, partial Fourier factor of 6/8, transverse slices, and phase encoding in the anterior-posterior direction. The last column of Table 1 reports the other parameters.

The SLM ghosting correction method presented here was compared to the vendor-provided ghosting correction solution based on linear phase correction. A one-dimensional navigator was played out at the beginning of each shot, consisting of three consecutive k-space lines acquired at the center of the k-space with positive, negative and positive readout polarities, respectively. The parameters of the linear model for correcting phase discrepancies between polarities were estimated from these calibration data (Buonocore & Gao, 1997). Corrected k-space datasets for each shot were interleaved to create a fully-sampled k-space dataset. After a 3D inverse Fourier transform, reconstructed coil-images were combined with the rSoS method (Roemer, Edelstein, Hayes, Souza, & Mueller, 1990).

Reconstructed images with both SLM and linear phase ghost correction approaches were then corrected for remaining geometric distortions induced by the diffusion-sensitizing gradients using the eddy tool in the FMRIB software library (FSL) toolbox (Andersson & Sotiropoulos, 2016).

To investigate the downstream effects of profound ghosting on the estimation of dMRI metrics, we carried out DTI with the linear least squares estimator, as implemented in the FSL tool ‘*dtifit*’ (Andersson & Sotiropoulos, 2016). Fiber ODFs were estimated with multi-shell multi-tissue constrained spherical deconvolution as implemented in the MRtrix3 software (Jeurissen & Tournier, 2014) (Tournier, et al., 2019). Response functions were estimated in white matter, gray matter, and CSF with the algorithm presented in (Dhollander, Mito, Raffelt, & Connelly, 2019). Default parameters were used.

## 3. Results

Axial diffusion-weighted images of the isotropic diffusion phantom acquired at *b* = 2 000 s/mm^2^ are presented in Fig 4. Visual inspection reveals noticeable residual ghosting artifact on the images reconstructed using linear phase correction and marked reduction of the ghosting when the SLM method is applied. Upon narrow windowing of the images, the ghosting artifacts are prominent both inside and outside the phantom, as best seen on the 10x scaled images. The ghosting artifacts are more pronounced for some diffusion directions, which is anticipated due to asymmetric eddy currents that may be more pronounced along certain gradient axes (e.g., G_z_ versus G_x_ or G_y_) (Setsompop, et al., 2013). However, the SLM-based method achieves a consistent reduction of ghosting for all diffusion directions, as quantified by the 10- to 20-fold reduction in ghost-to-signal-ratio (GSR) compared to the linear phase correction.

**Fig 4.**
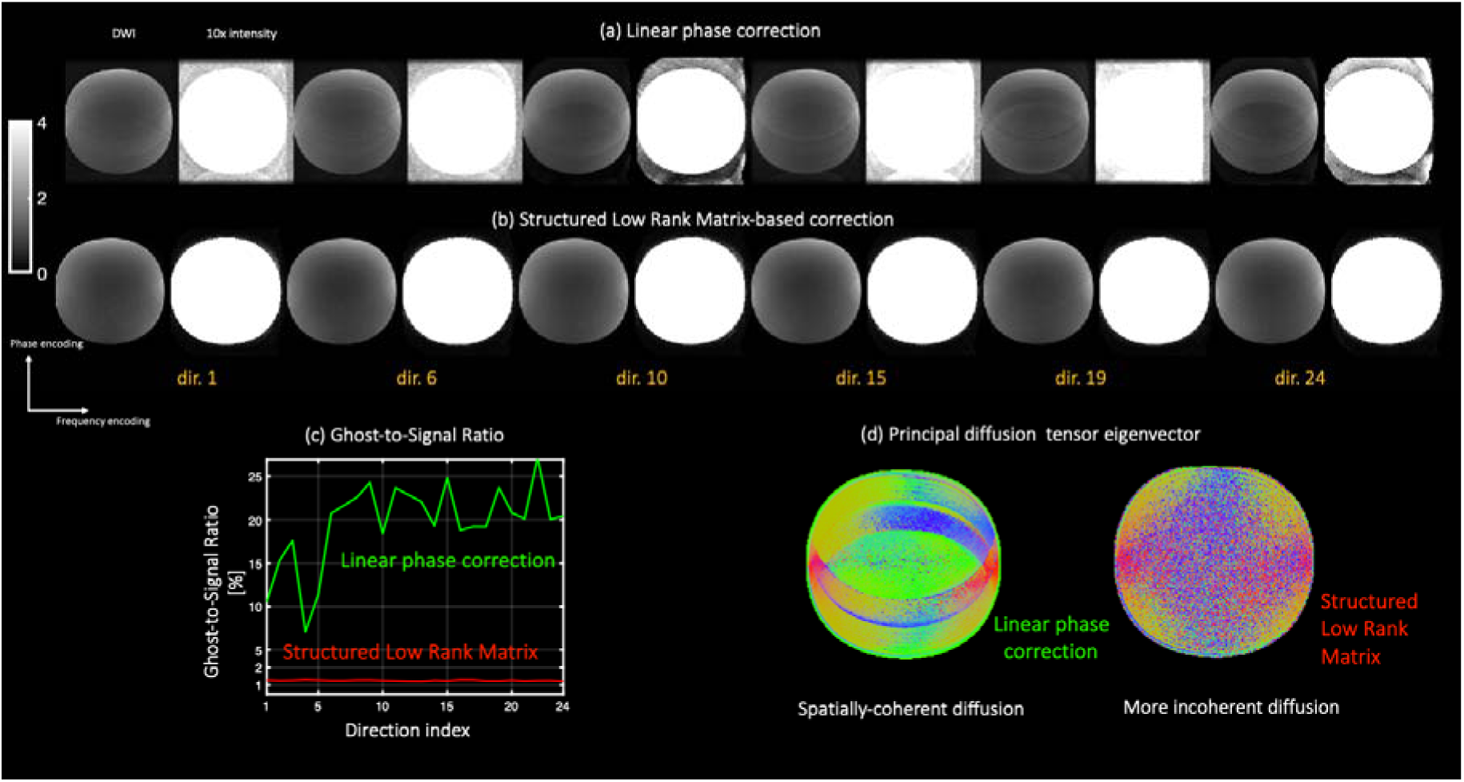
Axial diffusion-weighted MR images of an isotropic diffusion phantom (40% PVP solution) acquired at 1 mm^3^ isotropic spatial resolution, b = 2 000 s / mm^2^ and 24 diffusion-encoding directions. Images reconstructed with conventional linear phase correction (a) and with structured low-rank matrix-based method (b). Ghosting is drastically reduced when the SLM is used (see the ghost-to-signal ratio in (c)). Ghosting creates orientational bias in the main diffusion tensor eigenvector in the PVP solution (d), despite being an isotropic diffusion phantom. When ghosting is reduced with the SLM method, the orientational bias is suppressed.

This metric quantifies the level of ghost-only signal with respect of ghosting-free signal. Although the lack of a ground-truth is a known issue in the EPI ghost-correction literature, the GSR metric can provide insight into the degree of ghosting reduction without the need for a ghost-free reference acquisition, which may be difficult to achieve for a given set of acquisition parameters. As such, the GSR has been used extensively in other work to quantify the degree of ghosting reduction (Reeder, Atalar, & E.R., 1997) (Poser, Barth, Goa, Deng, & Stenger, 2013). The mean of the ghost-only signal is calculated in the area outside of the spherical phantom but limited by the coil sensitivity maps. While in the linear phase correction method the ghosting extends beyond this region, we restricted the calculation of GSR to within the coil-sensitivity profile map to provide a fair comparison with the SLM-method.

The SLM-method achieves GSR values of 1.5% on average for all 24 diffusion directions, whereas linear phase correction achieves much higher mean GSR values of 19.7% (Fig 4c). The lower value of the GSR with SLM-method indicates a superior ghosting reduction.

This experiment demonstrates that ghosting artifacts contaminate the dMRI signal, resulting in artifactual spatially coherent diffusion that is not present in the imaged sample. To demonstrate this effect, after reconstructing the images using linear phase correction and the SLM method, we estimated the diffusion tensor and displayed the primary eigenvector in Fig 4d. In an isotropic diffusion phantom, one would expect free diffusion without orientational preference, i.e., the directionality of the primary eigenvector should vary randomly across pixels, as the variability of the dMRI signal comes from noise only. Any coherent configuration can be attributed to artifactual signals contaminating the underlying dMRI signal. The primary diffusion eigenvector map obtained after linear correction clearly shows a preferred diffusion direction, which is attributed to the presence of coherent ghosts interfering with the accurate estimation of the underlying diffusion tensor. With the reduction of ghosting artifacts in the SLM-reconstructed images, the orientational bias of the primary diffusion eigenvector map decreases to the point where no preferred diffusion orientation is seen in most of the phantom.

Next, we compared the degree of ghosting reduction achieved with conventional linear phase correction compared to the SLM method in the ex vivo whole human brain specimen. Fig 5 shows representative axial, coronal, and sagittal views of the mean DWI averaged over all diffusion-encoding directions from the single-shell dMRI dataset (b = 4 000 s/mm^2^, ex-vivo dataset 1). Note the marked reduction in ghosting artifact seen in the images reconstructed with the SLM method compared to traditional linear-phase correction. The ghosting artifact is particularly pronounced near the ventricles on the coronal and sagittal views of the linear phase-corrected images (Fig 5c). The intensity variations attributed to ghosting are estimated to be up to 40% of the dMRI signal.

**Fig 5.**
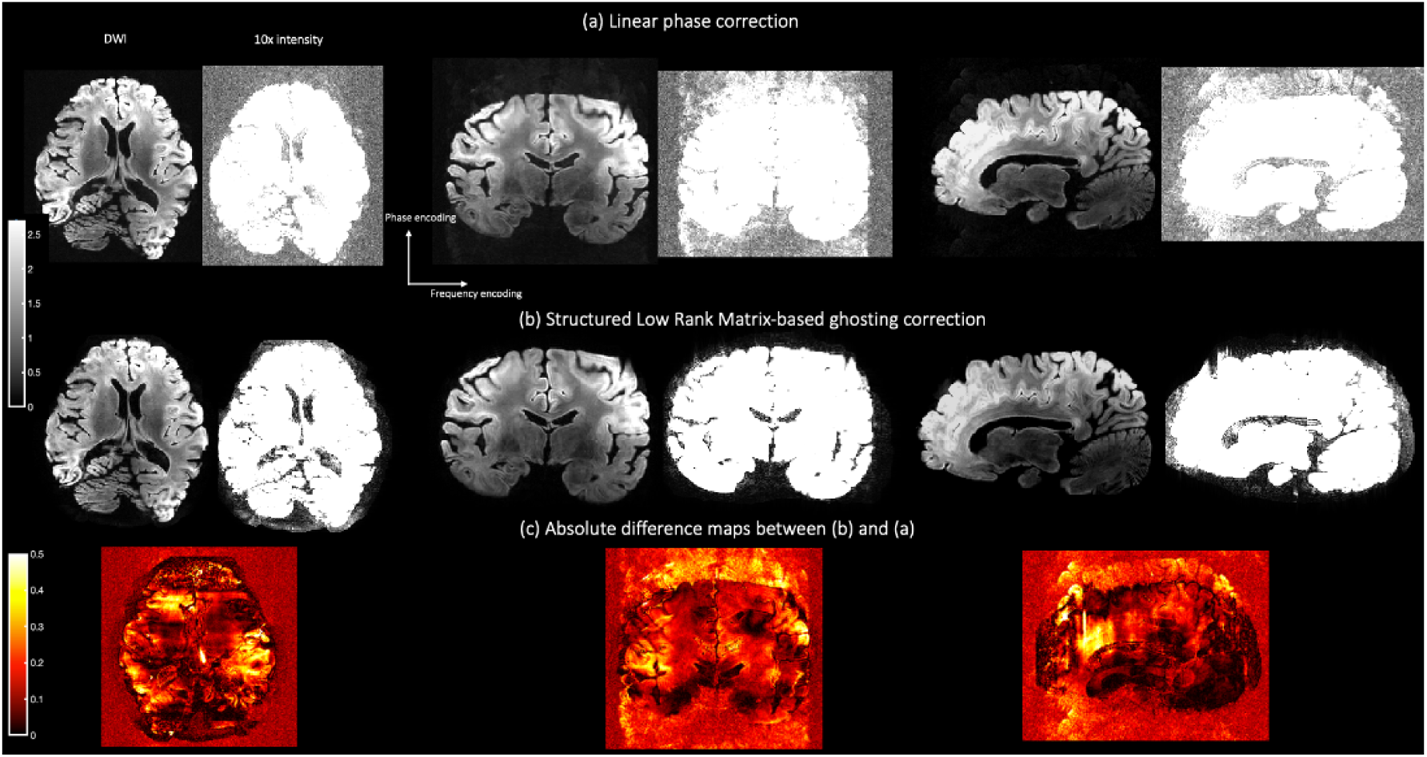
Axial, coronal, and sagittal views of the mean diffusion weighted data from ex-vivo dataset 1 averaged over all diffusion-encoding directions (0.8 mm^3^ spatial resolution at b = 4 000 s /mm^2^) reconstructed with (a) linear phase correction and (b) the proposed structured low rank matrix-based method. Note the marked ghosting reduction that is achieved with the SLM in comparison to the conventional linear-phase correction method. Scaled images multiplied by with a factor of ten are shown to emphasize ghosting suppression. Absolute difference maps between (b) and (a) illustrate the residual ghosting that is still present when linear phase correction is applied. The ghosting effect manifests as replication of ventricles in both coronal and sagittal slices.

Despite the known variability in ghosting severity across diffusion directions, the overall reduction in ghosting with SLM appears constant when averaged over the entire set of diffusion-encoding directions, which is not the case when linear phase correction is applied (Fig 6a). Ghosting artifacts are particularly noticeable in the cortex (green arrow in Fig 6c).

**Fig 6.**
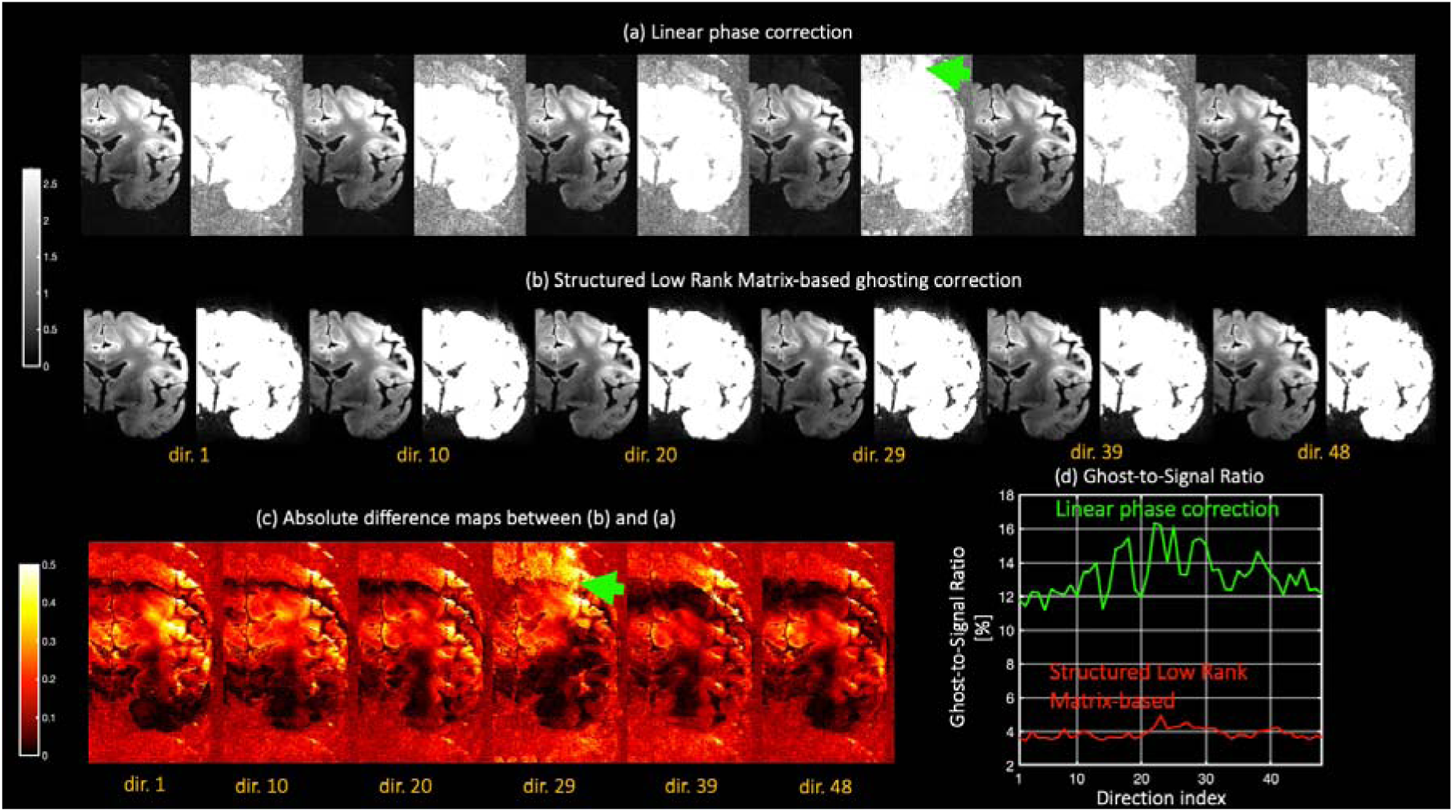
Reconstructed mean DWI’s from ex-vivo dataset 1 using (a) conventional linear-phase correction and (b) the structured low rank matrix-based method. A random selection of diffusion-encoding directions is presented. Note that ghosting is more intense for certain diffusion directions (compare direction 1 with direction 29, especially in cortex areas: green arrow) and is substantially reduced with SLM. The ghost-to-signal-ratio for this slice remains approximately constant for SLM and is more than that three times lower than that obtained with linear phase correction.

As with the phantom experiment, we calculated the GSR for the entire reconstructed volume. The mean of the ‘ghost signal’ was calculated in a region that excludes white and gray matter but includes CSF and extracranial structures that are contained in the spatial support of the estimated coil sensitivities. A GSR reduction of at least threefold was obtained when images were reconstructed with the SLM-based approach compared to conventional linear phase correction (Fig 7).

**Fig 7.**
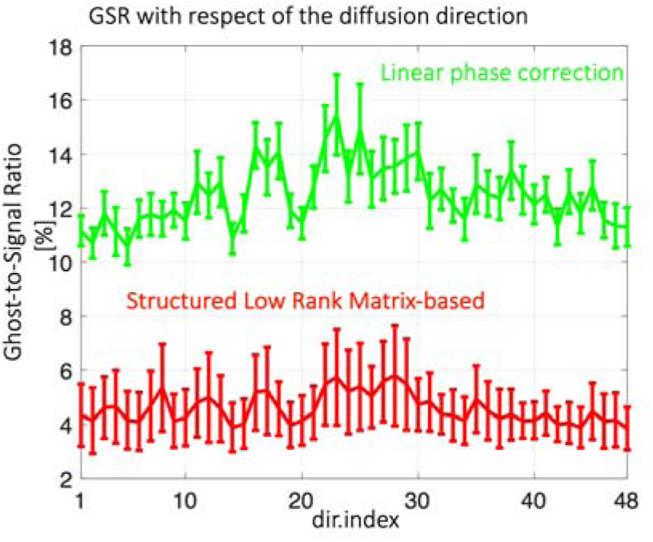
Ghost-to-Signal ratio depends on the index of the diffusion directions. Values are averaged over all slices. Error bars represent standard deviation.

Fig 8 shows coronal views of fractional anisotropy (FA) and mean diffusivity (MD) maps for two different slices reconstructed using linear phase correction and the SLM approach.

**Fig 8.**
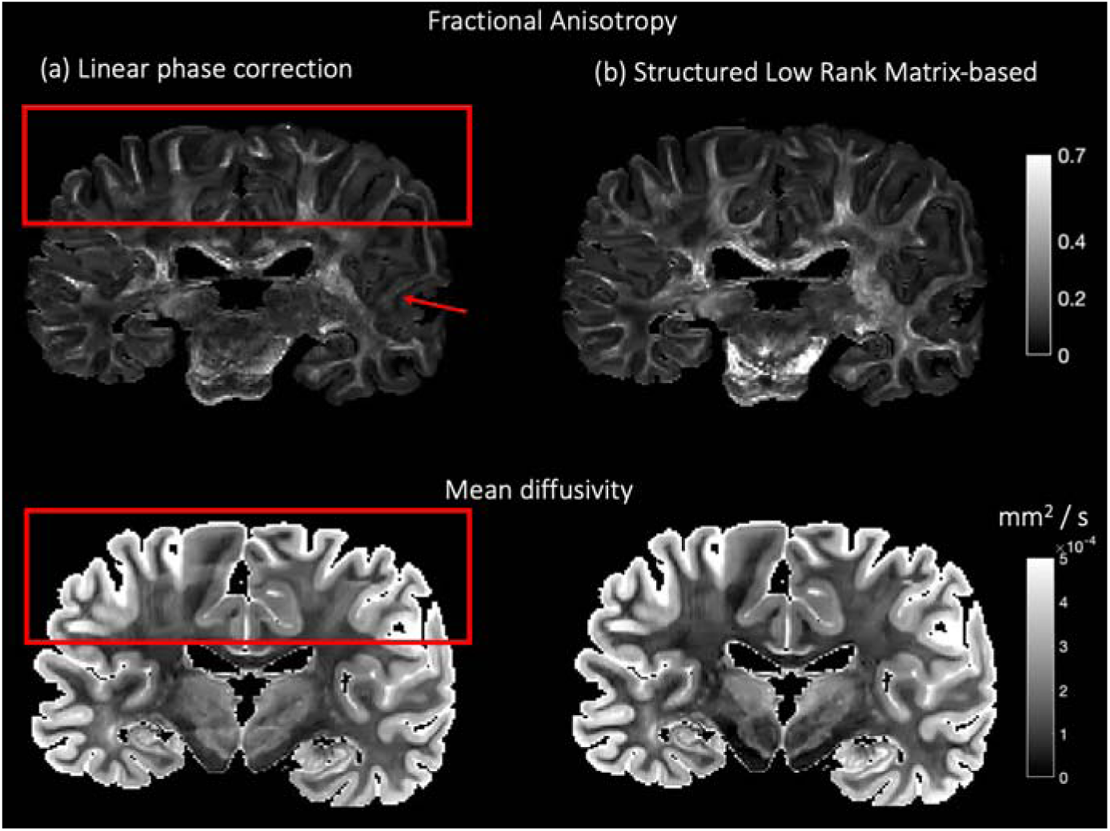
Fractional anisotropy (FA) and mean diffusivity (MD) maps from an ex-vivo DWI dataset obtained at b = 4 000 s/mm^2^ and 0.8 mm of isotropic spatial resolution (ex-vivo dataset 1), and reconstructed with a) conventional linear phase correction and b) structured low rank matrix-based method. Observe the prominent artifacts in the cortex on the FA and MD maps calculated from the linear phase-corrected data, and the missing tract in the FA map (see top, red arrow in the temporal lobe).

It is evident that residual ghosting artifacts on the linear phase-corrected images have a substantial impact on the calculated FA and MD maps. Artifactual replication of the cortical surface is clearly seen at the top of the coronal slices affecting the estimation of DTI measures in the cortex. Furthermore, discontinuities in the estimated FA can be seen in areas where fibers tracts terminate on the cortex, as well as abrupt and heterogeneous MD variations which do not correspond to distinct tissue configurations. Such artifacts are substantially reduced with the SLM method. As the ghosting is more intense in the temporal lobes (see absolute difference maps in Fig 5c or Fig 6c), it is not unusual to find missing white matter tracts in the FA maps derived from the linear phase-corrected data (red arrow in the temporal lobe), while the FA of the same tracts appear preserved when the ghosting is reduced with the SLM-based approach.

In addition to artifacts plaguing scalar metrics like FA or MD, ghosting also biases the diffusion orientation information. As shown in Fig 9, the principal eigenvector of the estimated diffusion tensor in the CSF reveals a coherent diffusion orientation of water, findings that were also observed in (McNab, et al., 2013) and were attributed to acquisition-related artifacts. This experiment (ex-vivo Dataset 1) and the previous phantom experiment provide further evidence that the orientational bias stems from residual ghosting. If, however, ghosting is reduced, diffusion in the CSF no longer remains coherent with defined anisotropy, as expected (see Fig. 9b).

**Fig 9.**
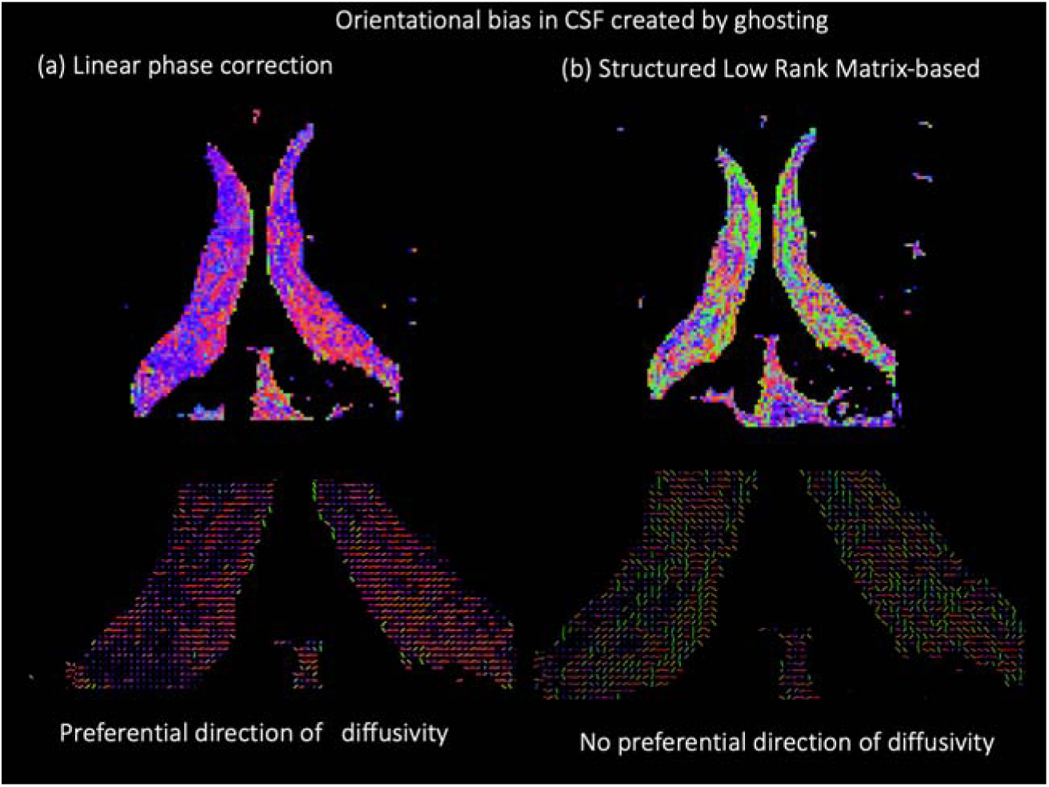
The orientational bias created by ghosting in the principal diffusion tensor eigenvector can easily be seen in isotropic-like areas as the cerebrospinal fluid (CSF) (a). When ghosting is substantially reduced, as with the SLM method, the principal diffusion tensor eigenvector does not show any preferential direction (b), as theoretically expected.

Orientational bias created by ghosting affects tissues beyond the CSF, e.g., white and gray matter. An example is given in Fig 10.

**Fig 10.**
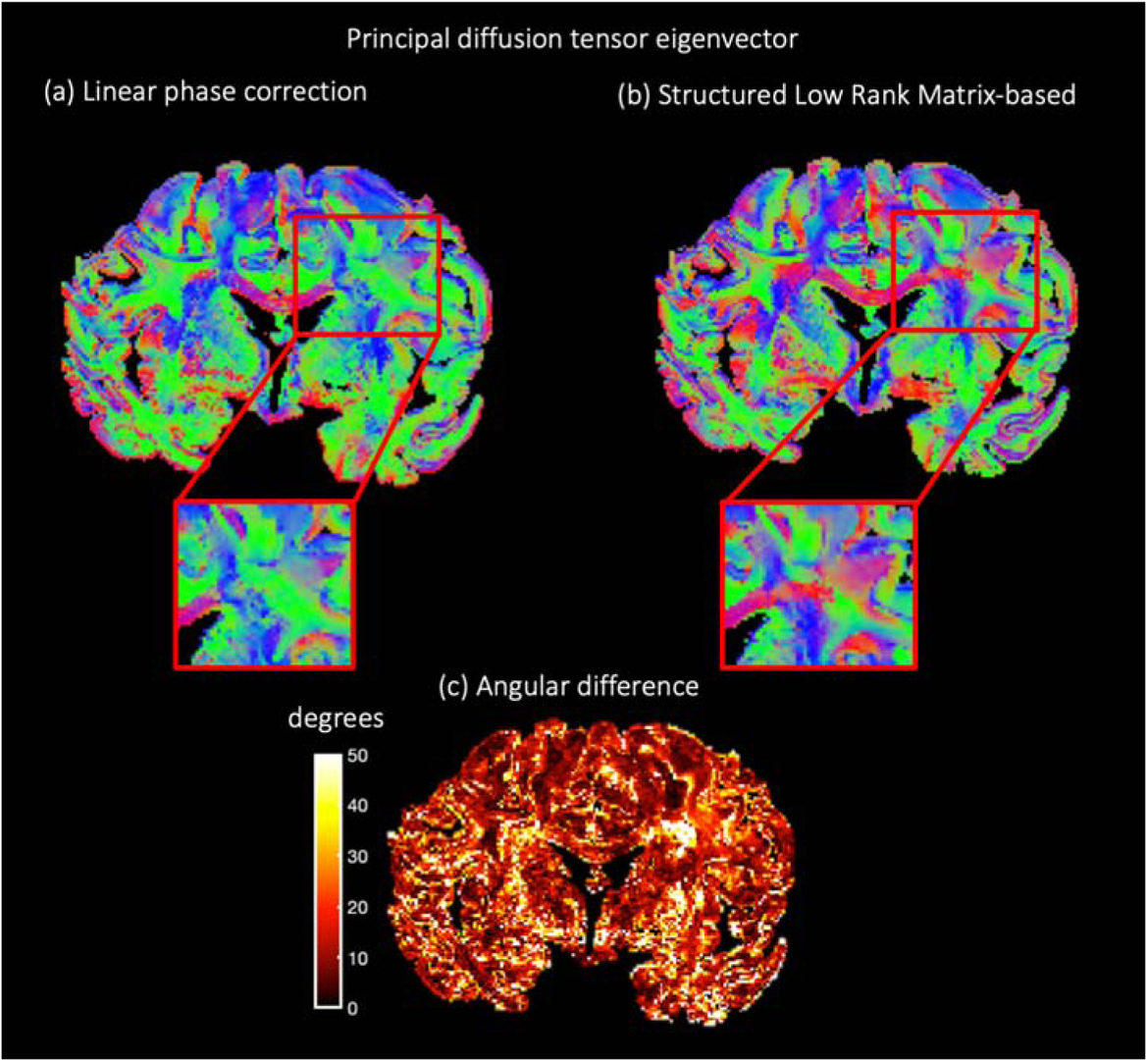
The orientational bias created by ghosting in the principal diffusion tensor eigenvector can easily be seen in isotropic-like areas as the cerebrospinal fluid (CSF) (a). When ghosting is substantially reduced, as with the SLM method, the principal diffusion tensor eigenvector does not show any preferential direction (b), as theoretically expected.

Fig 10 shows a coronal view of the principal diffusion tensor eigenvector map. Ghosting-induced orientational bias alters directionality information, introducing an orientation bias along the ventral-dorsal direction (green color), while directionally information along left and right (red color) seems to be lost. A decrease in diffusivity along left-right direction can be observed in cortical areas and especially in the corona radiata region, which is magnified for a better visualization. Though the ground-truth is currently inaccessible here, it is revealing to report the angular differences between the main diffusion eigenvector after ghosting correction with the SLM-based approach and with the linear phase correction. Angular differences values are high especially in areas where crossing fibers are known to exist. Along this line, the experiment with fiber ODFs in the ex-vivo multi-shell data provide further evidence to the strong effect of residual ghosting in mapping complex fiber configurations.

To further emphasize the benefit of the proposed methodology for accurate submillimeter whole ex-vivo human brain dMRI with high-gradient strengths, we present results showing the impact of unwanted ghosting in neuroanatomical areas distributed throughout the whole human brain, including the pons, cerebral cortex, and corpus callosum (Fig 11a). Insufficient reduction of the ghosting hampers reliable mapping of the transverse pontine fibers (Fig 11a) and intracortical diffusivity (Fig 11b). When ghosting is minimized with the advanced reconstruction approach presented here, a much better delineation of pontine fibers can be obtained, and radial diffusivity in cortical areas can be recovered (magnified region in Fig 11b). Corpus callosum fibers obtained after applying the SLM-method show less angular dispersion than those obtained after conventional linear phase correction.

**Fig 11.**
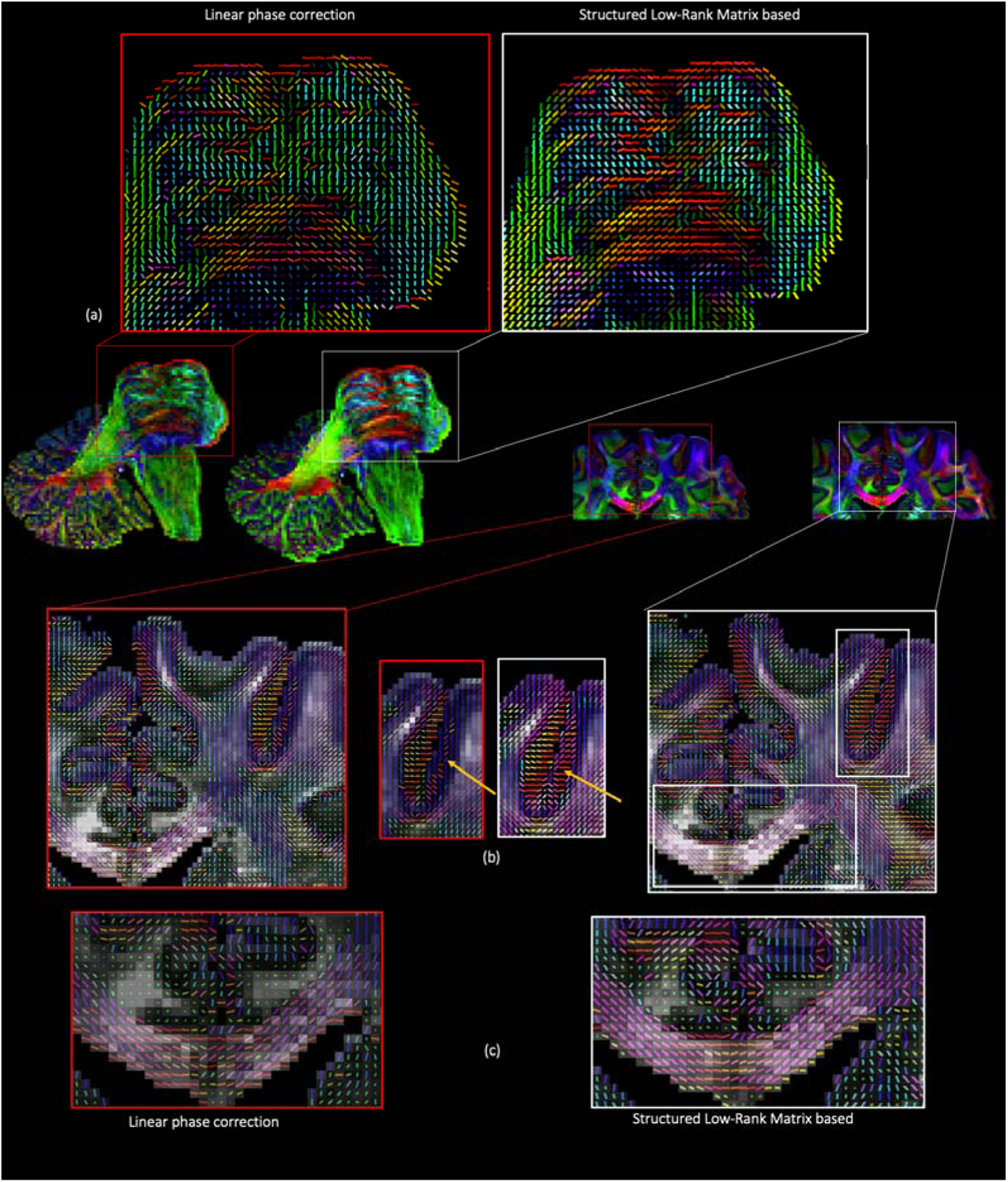
Several neuroanatomical regions impacted by insufficient ghosting correction and better resolved with structured low-rank matrix based EPI correction. See the better delineation of pontine fibers in the brain pons (a) as well the radial diffusivity pattern in the cortex, which is obscured when using conventional linear-phase methods for ghosting correction. Commissural fibers along corpus callosum look more coherent with SLM based in comparison to that estimated with conventional linear phase correction technique.

Results from the multi-shell dataset (ex-vivo dataset 2) are presented in Fig 12, Fig 13 and Fig 14. A coronal slice of two diffusion-weighted images acquired with *b* = 4 000 and 10 000 s/mm^2^ is shown in Fig 12. In agreement with previous experiments, images reconstructed with the proposed SLM-based reconstruction shows substantially superior ghosting elimination compared to images corrected with linear phase correction. Notice the pronounced ghosting reduction in the CSF, especially in the higher shell. In this case, the gradient strength was close to the nominal limit of the 3T Connectom scanner (300 mT/m).

**Fig 12.**
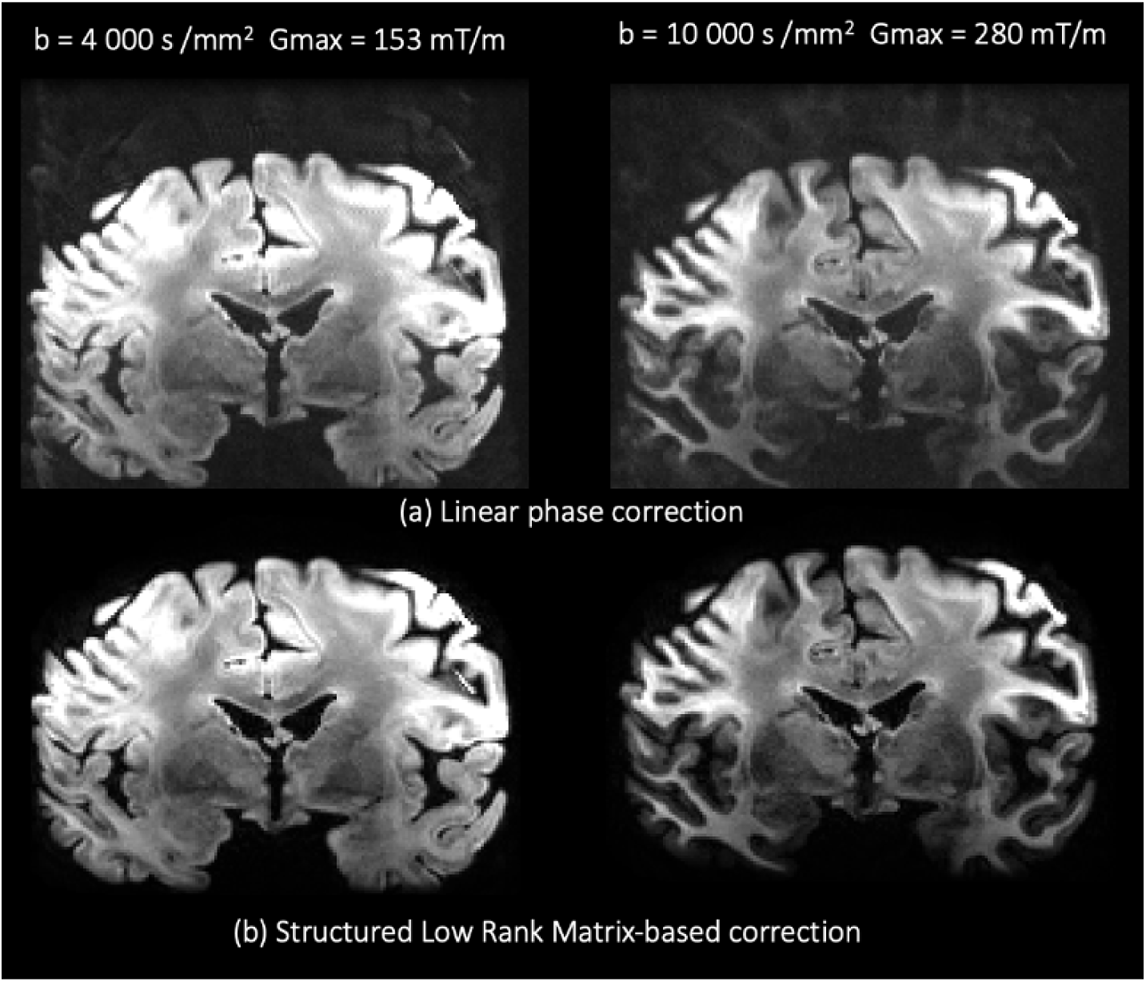
Diffusion weighted images from the experiment with multi-shell data, (0.8 mm^3^ at b = 4 000 and 10 000 s/mm^2^) with (a) ghosting elimination with linear phase correction and (b) the proposed Structured Low Rank Matrix-based method.

**Fig 13.**
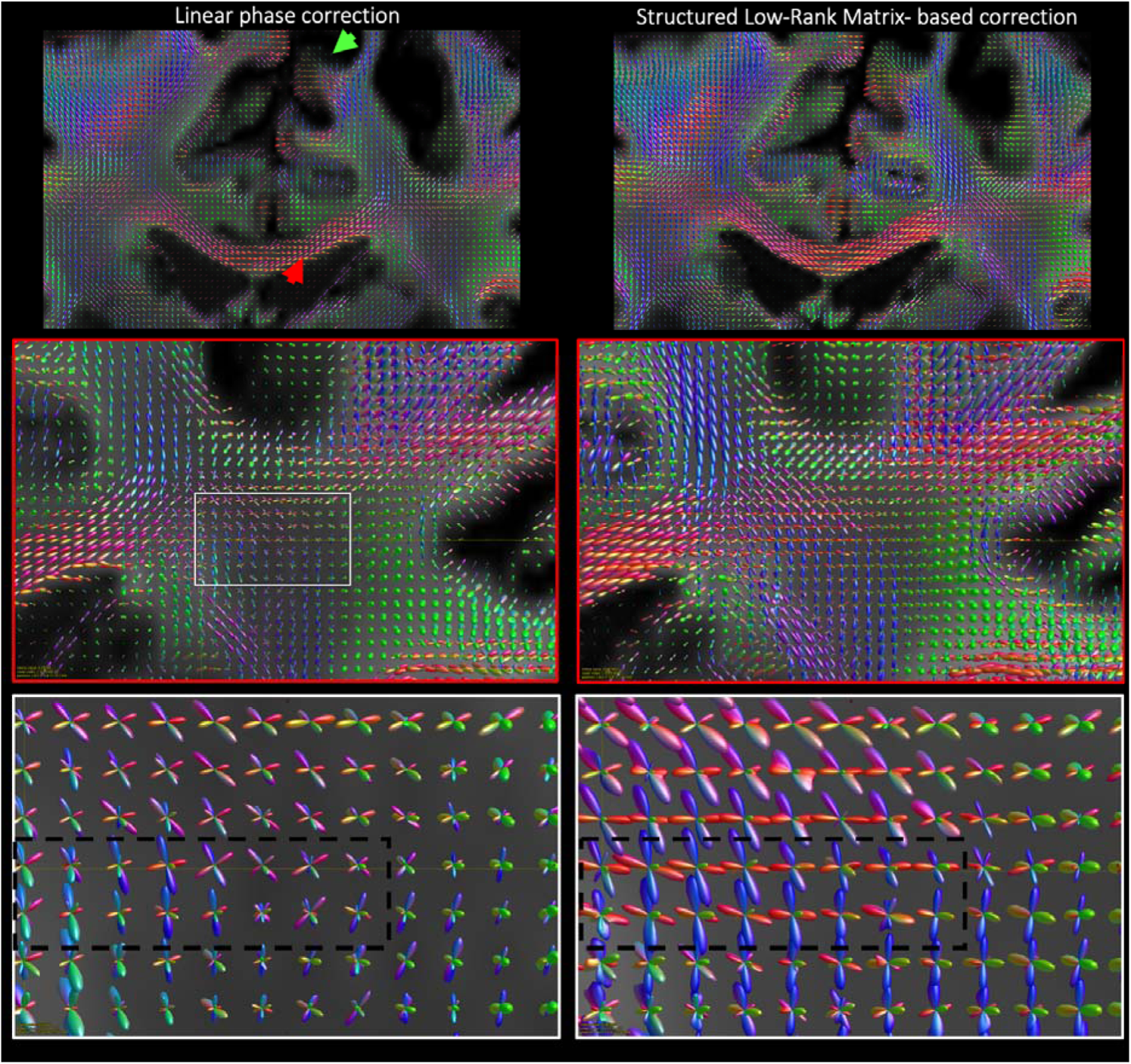
Fiber ODFS estimated with multi-shell multi-tissue spherical deconvolution from the multi-shell dMRI dataset ghosting-corrected with (a) linear phase correction and (b) the Structured Low Rank-Matrix based methodology. There exists a consistent decrease in the amplitude of the fODFS peaks when technique (a) is applied (see green arrow). Note as well that remaining ghosting introduces higher angular dispersion the fODFs, see magnified areas in the corona radiata or in the corpus callosum (red arrow). More coherent orientation can be found after reducing ghosting with the Structured Low Rank-Matrix based method.

**Fig 14.**
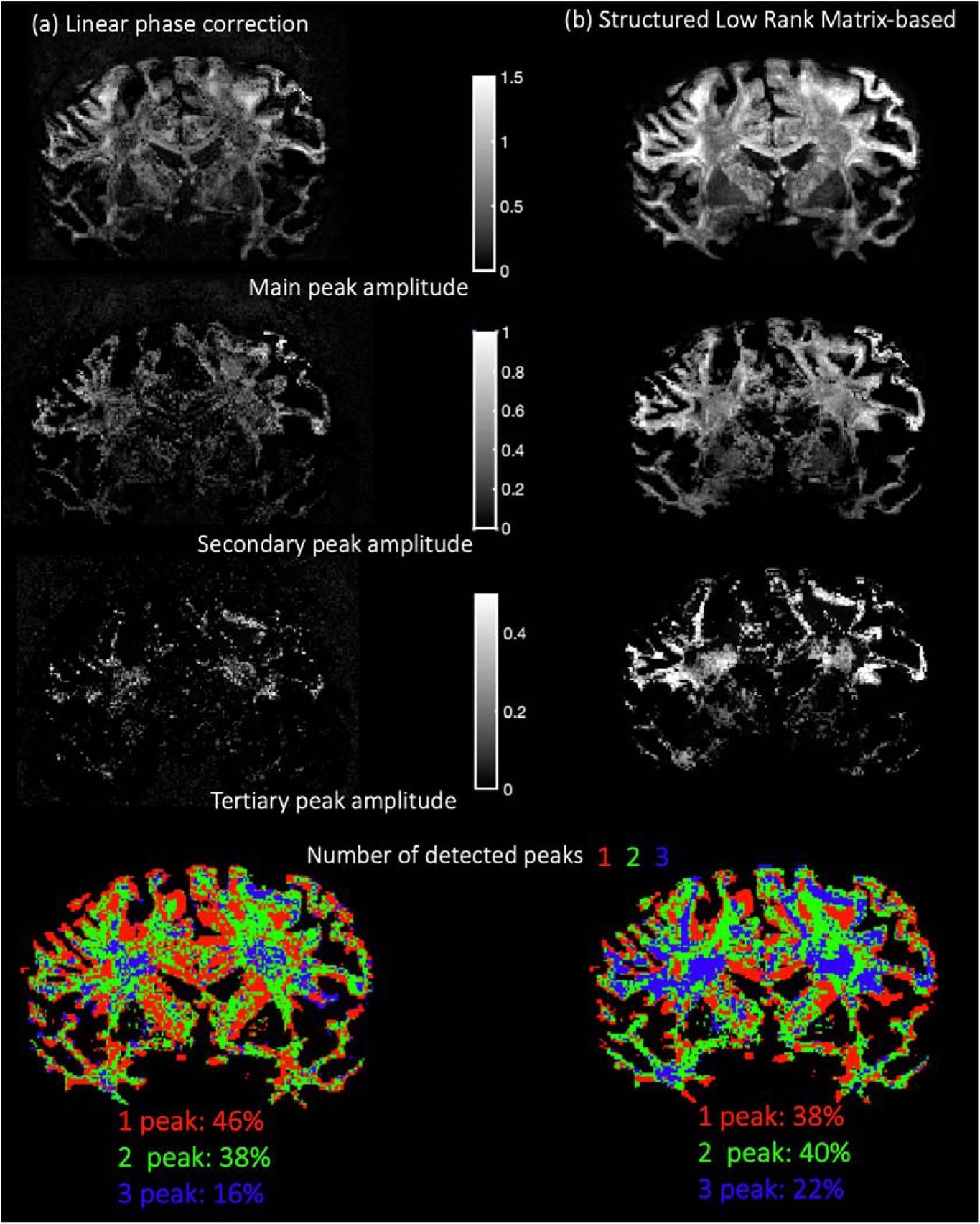
Amplitude of the main, secondary, and tertiary peaks of the fiber ODFs estimated after ghosting correction with (a) linear phase correction and (b) Structure Low Rank Matrix-based reconstruction. The number of detected peaks is also shown. Note the overall increase in amplitude of the peaks maps when ghosting is further reduced with the SLM method as well as the higher detection rate of complex-fiber configurations.

Fiber ODF maps are presented in Fig 13. An immediate observation is that amplitudes of the fiber ODFs are consistently larger when ghosting is reduced with the SLM-based approach than when linear phase correction approach is applied (for example, see the cortical region marked with a green arrow). In general, fODFs show less dispersion and present a more coherent spatial pattern, notably in a crossing-fiber area (magnified image in Fig 13) and in the corpus callosum (see red arrow). Note as well the consistent finding of 3-peak fODFS that are displayed in the magnified, dashed black rectangle, where corpus callosum projections, cortical spinal tract, and superior longitudinal fasciculus are known to intersect each other.

Crossing fibers in the corona radiata estimated after ghosting reduction with linear phase correction present less spatially coherent angular orientation. We quantified the amplitude of the primary, secondary, and tertiary peaks of the fODFs. Corresponding maps are displayed in Fig 14. Note the consistent decrease in intensity for the three maps and the spatial variability across voxels when ghosting artifacts are not diminished enough.

The number of detected peaks is displayed in Fig 14 as well. More complex fiber configurations can be detected when ghosting is suppressed with the SLM-based method. Indeed, the percentage of complex-fiber configurations (>1 peaks) (Jeurissen, et al., 2013) (Fan, et al., 2014) was found to be 62 % in contrast to 54 % when linear phase correction was applied. Note as well that in the corona radiata, the number of detected peaks follows a smoother spatial pattern rather than the more arbitrary configurations observed with linear phase correction.

## 4. Discussion

In this paper, we have shown that ghosting is a pervasive artifact that may bias the estimation of dMRI measures derived from ultra-high spatial resolution ex vivo diffusion MRI using strong gradients on human MRI scanners. We have demonstrated that when using 3D multi-shot EPI sequences targeting high SNR, ghosting may not be sufficiently mitigated with typical vendor solutions (linear phase correction with one dimensional navigator). Ghosting not only creates artifactual effects that contaminate the diffusion weighted images but also biases subsequent diffusion analyses. Conventional EPI-techniques usually assume simplistic models for the phase differences between polarities and shots (Buonocore & Gao, 1997) (Chen & Wyrwicz, 2004) (Hennel, 1998). When using strong diffusion gradients, as shown in this paper with a 3T Connectom scanner, the assumption of linear phase does not hold. The strong eddy currents generated by the diffusion-encoding gradients introduce non-linear variations in the background magnetic field that modulate the phases of different k-space lines and shots. We have proposed an advanced reconstruction method that corrects EPI-ghosting with SLM-based methods. This approach does not require additional scans and treats the EPI-ghosting correction step as a unified, under-sampled k-space reconstruction problem. The k-space dataset is divided into subsets, each one comprising k-space lines acquired at a given excitation/shot and readout gradient polarity. A ghost-free image is reconstructed from the undersampled k-space data of each of these subsets. To solve this ill-posed reconstruction problem, constraints based on SLM modeling are enforced on the reconstructed images. Those constraints leverage the fact that the k-space of the reconstructed image is linearly predictable as the images are highly correlated.

We showed that this advanced reconstruction approach was able to reduce ghosting artifacts substantially. The SLM-based EPI correction method was first validated in an isotropic diffusion phantom and then in fixed whole human brain that were acquired at 0.8 mm isotropic resolution and *b*-values up to 10 000 s/mm^2^. Reliable DTI measures and fiber orientation distribution functions (fODFs) were recovered from the reconstructed images. We compared the residual ghosting artifacts from SLM-based EPI correction and linear phase correction to illustrate the substantial improvement in visualizing key neuroanatomical areas distributed across the whole brain, enabling detailed dMRI mapping of such diverse structures as the central pontine fibers, cortical anisotropy, and fibers in the corpus callosum, respectively.

In particular, the proposed technique based on SLM modeling achieved superior ghosting reduction, assessed both visually and quantitatively, and leads to more reliable diffusion analysis. Maps of scalar dMRI metrics such as FA and MD, which were plagued by ghosting artifacts with conventional linear phase correction, were substantially less affected by this pervasive artifact with SLM reconstruction.

As the ghosting varies depending on the direction and amplitude of the diffusion-encoding gradients, it perturbs the dMRI signal in such a way that it introduces an orientational bias in directional diffusion metrics. We have confirmed the existence of orientational bias first with an isotropic diffusion phantom, where coherent anisotropy was observed due to insufficient ghost correction. Orientational biases in the estimation of DTI metrics in the ventricles of ex vivo whole human brain samples using strong diffusion-encoding gradients have been also reported in other studies (McNab, et al., 2013). Our experiments agree with this observation and point to ghosting may be the main underlying cause. Indeed, when ex vivo dMRI datasets are reconstructed with the methodology proposed here, diffusion in the ventricles did not show any preferential direction.

The orientational bias clearly affects both white and gray matter. We have shown that insufficient ghosting reduction introduces an artifactual diffusion orientation in distributed neuroanatomical regions of importance, such as the brainstem, cerebral cortex, or large white matter fiber bundles. Considerably better delineation of the cerebellopontine fibers in the pons is achieved when ghosting is reduced with the SLM-based method presented here. Fibers in the corpus callosum are also shown to follow a more coherent organization.

Results with high-angular diffusion MRI revealed that the estimated fiber ODFs are severely contaminated with the pervasive ghosting artifact. Nevertheless, if ghosting is reduced with the SLM-based technique, the fODFs follow a smoother, less arbitrary spatial distribution across voxels. Besides reduced angular dispersion, a higher percentage of complex fibers configuration, especially three-crossing fibers, can be resolved when ghosting is accounted for with SLM modeling.

Although not included in this initial study, we anticipate that the proposed technique will have a positive impact in improving brain connectivity analyses that are based on tractography-based approaches. Similarly, accurate cortical laminar diffusion analysis is likely to necessitate advanced EPI-ghosting methodologies like the one presented here. With a moderate *b*-value of 4 000 s / mm^2^ and high gradient strength of 180 mT/m, we have shown that intracortical diffusivity deteriorates in the areas of cortex where the ghosting artifacts are more prominent. With the SLM-based ghosting correction approach, we recovered radial diffusivity along the surface of cortical regions that were otherwise greatly affected by the ghosting artifact. We envision that the use of such sophisticated EPI correction approaches as SLM will become even more important and necessary when resorting to ultra-high *b*-values on dedicated high-gradient human MRI scanners (Fan, et al., 2016). Those studies are part of our future work.

We would like to mention some limitations of the current work and possible future extensions. The computation time of our relatively optimized implementation of the SLM-method presented here is long, around an hour for each slice of the ex-vivo dataset. The implementation is done in MATLAB on a computer with an Intel Xeon Gold 2.7GHz (28 cores) processor and 512 GB of RAM. We expect that code migration to other programming languages as C and/or the application of dimensionality reduction techniques such as array compression approaches (Buehrer, Pruesmann, Boesiger, & Kozerke, 2007) would shorten this computation time considerably. Though the proposed method achieved more than three-fold ghosting reduction as measured with the ghost-to-signal ratio, ghosting artifacts may remain and contaminate the dMRI signal. In this sense, simple improvements to the SLM-based algorithm, for example, tuning the regularization parameter λ and the matrix rank Y per every slice direction can probably reduce ghosting even further. Adaptation of the proposed methodology to a three-dimensional reconstruction is, in principle, conceptually straightforward and can leverage linear k-space relationships between adjacent slices. Nevertheless, we expect this approach to require prohibitively large RAM. Finally, it should be noted that other more advanced SLM methods that use additional scan(s) for ghosting reduction could help reducing the artifact further (Lobos, et al., 2021), although a detailed investigation and comparison of different SLM-based methods is beyond the scope of this work.

Currently, the k-space data for each diffusion direction are reconstructed separately. The fact that ghosting varies across diffusion direction suggests that it may be advisable to reconstruct the whole set of dMRI images simultaneously by exploiting similarities of k-space acquired at different diffusion directions or q-space values. Those approaches are becoming increasingly popular in the dMRI literature MRI (Ramos-Llordén, et al., 2020) (Wu, Koopmans, Andersson, & Miller, 2019) (Mani, Magnotta, & Jacob, 2021) (Hu, et al., 2020) (Ramos-Llordén, Aja-Fernández, Liao, Setsompop, & Rathi, 2019) (Haldar, et al., 2013) (Wu, Koopmans, Andersson, & Miller, 2019) (Wu, et al., 2014). Exploratory analyses along these lines are an important aspect of our future work.

The use of field-monitoring approaches for eddy current-induced artifacts deserve special mention (Wilm, Barmet, Pavan, & Pruessmann, 2011). In section 2.1, the possibility of combining the technique presented here with this hardware solution was outlined. Field-monitoring approaches necessitate additional hardware, a dynamic field camera, but they can provide additional benefits such as mapping the temporal B0 field variations that interfere with the nominal k-space trajectory. As such, on top of ghosting correction, remaining geometrical distortions and the induced spatial misalignment between DWI are accounted for in a single step. To better condition the inverse reconstruction problem, constraints based on SLM modeling theory, as those we have used in this work, can be easily accommodated.

Though a comprehensive validation is beyond the scope of the paper, we believe the proposed method may be useful for in-vivo EPI dMRI with strong diffusion strengths as well, since exacerbated eddy currents are equally present in this case at the high strength gradient regime.

## 5. Conclusion

The field of neuroanatomy can greatly benefit from ex-vivo diffusion MRI to resolve structures at mesoscopic (sub-mm) scale that are inaccessible with in vivo dMRI due to limited SNR. To achieve ultra-high-resolution, high b-value dMRI with short image readouts, and with large FOV, one can take advantage of ultra-strong diffusion gradients such as those available on the Siemens 3T Connectom scanner. At this high gradient-strength regime, the gradient-induced eddy currents become a significant issue. The strong EPI-ghosting artifacts in multi-shot DWI-EPI sequences, one of the common pulse sequences in ex-vivo dMRI, contaminate the reconstructed images substantially. We demonstrated here that conventional vendor solutions for ghosting mitigation based on linear-phase correction methods fall short, confounding subsequent diffusion analysis. We then proposed an advanced image reconstruction methodology that uses SLM-based methods for EPI correction, showing superior ghosting reduction thereby enabling more reliable diffusion mapping of ghosting-affected areas such as the human brain pons, cortex, corpus callosum or corona radiata.

The proposed method was validated in ex vivo dMRI experiments performed on a whole human brain, acquired at 0.8 mm isotropic resolution with *b*-values and maximum gradient strengths up to 10 000 s/mm^2^ and 280 mT/m, respectively. An isotropic diffusion phantom was also used to confirm the validity of the technique in a controlled scenario. Supported by our experimental validation, we advocate for the use of specialized reconstruction techniques such as the one presented here to achieve accurate and reliable submillimeter whole-brain ex-vivo diffusion MRI with strong diffusion gradients.

## Acknowledgements

This work was supported by the National Institutes of Health under grant numbers P41-EB015896, P41-EB030006, U01-EB026996, R01-NS118187, R01-EB021265, R03-EB031175, 01-EB028797, U01-EB025162, and K23-NS096056.

## Abbreviations

48ch: 48-channel
CSF: Cerebrospinal Fluid
dMRI: Diffusion Magnetic Resonance Imaging
DTI: Diffusion Tensor Imaging
DWI: Diffusion-Weighted Image
FA: Fractional Anisotropy
fODF: fiber Orientation Distribution Function
GSR: Ghost-to-Signal ratio
MD: Mean Diffusivity
MSMT-CSD: Multi-Shell Multi-Tissue Constrained Spherical Deconvolution
MM: Majorize-Minimization
PVP: Polyvinylpyrrolidone
ss: single-shot
SLM: Structured Low-rank Matrix
SNR: Signal-to-Noise ratio
TE: Echo time
TR: repetition time

